# Guide RNA binding induces reverse transcription bias during RT-qPCR analysis of RNA-targeting CRISPR/Cas systems

**DOI:** 10.64898/2026.08.16.745121

**Authors:** Boyu Huang, Carlos Orosco, Elijah Stewart, Mahika Balaraju, Yasmin B. Elhabashy, Piyush K. Jain

**Affiliations:** Department of Chemical Engineering, University of Florida, Gainesville, FL, USA; Department of Molecular Genetics and Microbiology, University of Florida, Gainesville, FL, USA; UF Health Cancer Center, University of Florida, Gainesville, FL, USA

## Abstract

RNA-targeting CRISPR systems are commonly evaluated by RT-qPCR, but guide RNA binding can confound these measurements. We show that crRNA alone produces apparent knockdown without reducing target RNA abundance, whereas RNA sequencing remains unbiased and reveals guide-associated transcriptomic perturbations. A simple RNA denaturation step before reverse transcription restores accurate RT-qPCR quantification, providing practical guidance for RNA-targeting CRISPR analysis and guide design.

## Main

RNA-targeting CRISPR/Cas systems have emerged as versatile tools for programmable gene silencing, enabling transient depletion of target transcripts without permanent genome editing^1-3^. Knockdown efficiency is routinely assessed by reverse transcription quantitative PCR (RT-qPCR)^4^, which is widely assumed to faithfully reflect RNA abundance. Although CRISPR RNAs (crRNAs) are designed to direct target recognition and cleavage^5, 6^, whether crRNA binding itself influences RT-qPCR measurements has remained unexplored. While benchmarking RNA-targeting CRISPR systems, we unexpectedly observed substantial apparent reduction in target RNA even in the absence of RNA-targeting nuclease activity. Here, we show that crRNA binding introduces a reverse transcription-dependent bias that distorts RT-qPCR measurements, whereas RNA sequencing remains unaffected. A simple RNA denaturation step before reverse transcription restores accurate quantification.

While evaluating evoCas7-11-mediated RNA knockdown, we included an unrelated AsCas12a nuclease as a negative control, as AsCas12a cannot recognize the evoCas7-11 crRNA or cleave the target RNA (Fig. 1a)^7-9^. Unexpectedly, co-transfection of the evoCas7-11 crRNA with AsCas12a produced an apparent reduction in target RNA comparable to that observed with the cognate evoCas7-11 nuclease. Because AsCas12a is incapable of mediating RNA cleavage in this experimental setting, RNA-guided target cleavage cannot account for the observed reduction in RT-qPCR signal. Instead, these findings suggested that the apparent reduction in target RNA could occur independently of the cognate Cas nuclease. We therefore sought to determine whether this unexpected activity was an intrinsic property of the crRNA itself.

**Fig 1:**
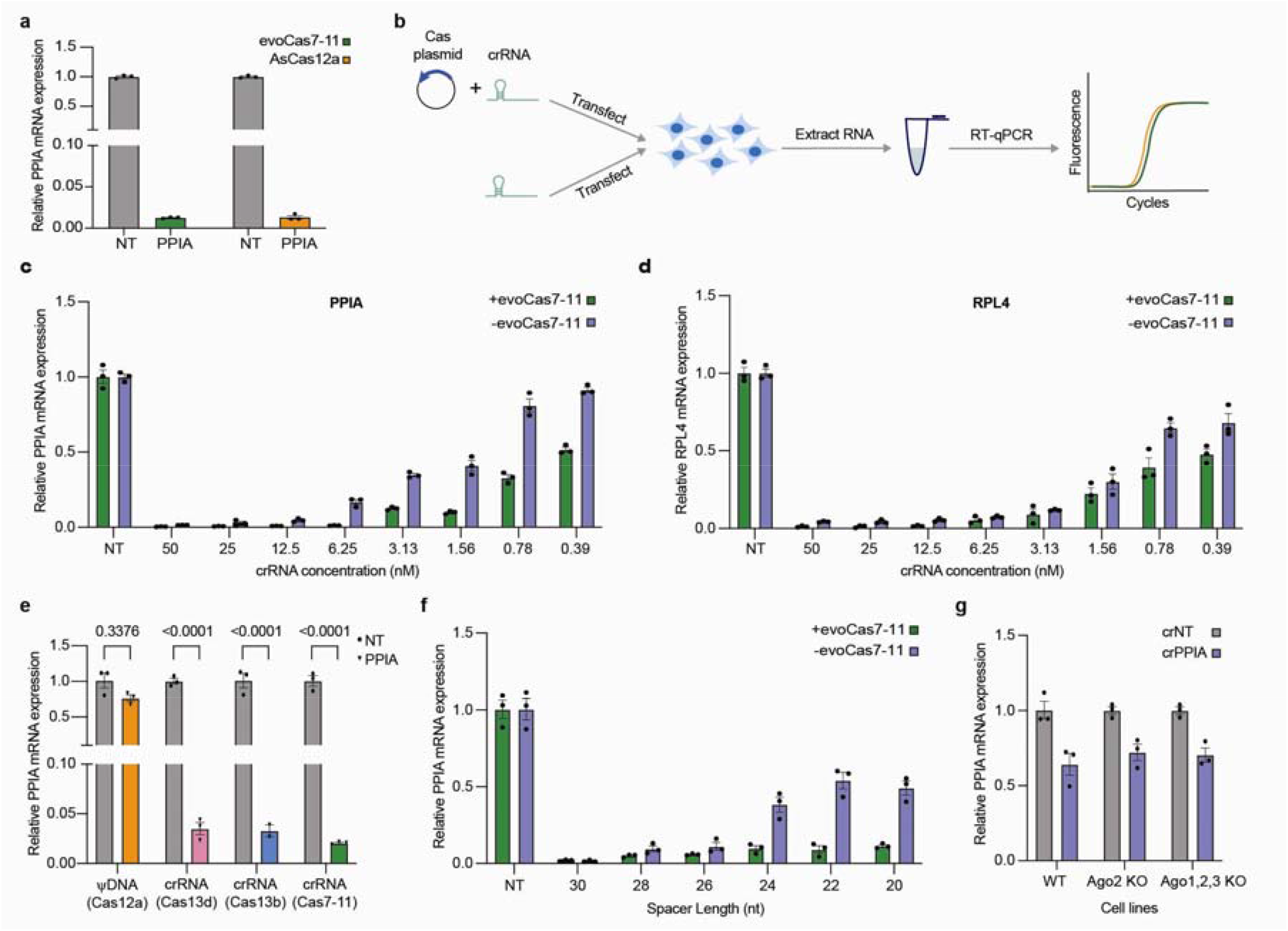
crRNAs alone induce apparent target depletion independently of cognate RNA-targeting nucleases. **(a)** RT-qPCR analysis of PPIA following co-transfection of evoCas7-11 crRNA with plasmids expressing either evoCas7-11 or the non-cognate nuclease AsCas12a. **(b)** Schematic of the experimental workflow for evaluating crRNA-dependent effects on RT-qPCR quantification. **(c, d)** Dose-dependent effect of evoCas7-11 crRNA on the apparent reduction of *PPIA* **(c)** and *RPL4* **(d)** measured by RT-qPCR. **(e)** Comparison of guide molecules with distinct architectures and chemistries targeting the identical RNA sequence. Statistical significance was determined by ordinary two-way ANOVA with Šídák’s multiple-comparisons test. **(f)** Effect of spacer truncation on apparent target RNA reduction measured by RT-qPCR. **(g)** RT-qPCR analysis in Ago-deficient cells. **(a, c-g)** Data are presented as mean ± s.e.m. from three biologically independent experiments (n = 3).

To further characterize this unexpected activity, we next investigated its dependence on crRNA abundance and architecture (Fig. 1b). For both PPIA and RPL4, the apparent reduction in target RNA increased with crRNA abundance and remained evident even at low guide concentrations, indicating that the phenomenon is not restricted to high crRNA inputs (Fig. 1c, d). We next asked whether this activity was unique to the evoCas7-11 guide architecture. Unexpectedly, guide RNAs from three distinct RNA-targeting CRISPR/Cas systems produced comparable apparent reduction in target RNA despite their different guide architectures, whereas our previously developed DNA guide (ΨDNA) targeting the identical RNA sequence had no detectable effect (Fig. 1e)^10-12^. Progressive shortening of the spacer markedly reduced the apparent reduction in target RNA, indicating that the activity depends on the extent of guide–target complementarity (Fig. 1f). Collectively, these data suggested that the apparent reduction in target RNA is driven by the RNA guide itself rather than the cognate RNA-targeting nuclease.

Disruption of the endogenous RNA interference pathway had no detectable effect on the apparent reduction in target RNA (Fig. 1g), excluding canonical Ago-mediated silencing^13,^ . ^14^These observations prompted us to investigate whether the apparent reduction in target RNA reflected genuine transcript depletion or instead arose during RNA quantification.

To distinguish between genuine transcript depletion and a quantification artifact, we established an RNA spike-in assay in which synthetic crRNAs were added directly to total RNA isolated from untreated cells immediately before reverse transcription (Fig. 2a). Addition of crRNAs alone was sufficient to reproduce the apparent reduction in target RNA observed in intact cells. Consistent with our cellular observations, the apparent reduction in target RNA increased with crRNA amount and progressively diminished as the spacer sequence was shortened (Fig. 2b, c). Because crRNAs were introduced only after RNA isolation, the observed reduction cannot result from RNA degradation or any cellular RNA silencing pathway. Instead, these findings indicate that the apparent reduction in target RNA is introduced during RNA quantification rather than reflecting a genuine change in RNA abundance.

**Fig 2:**
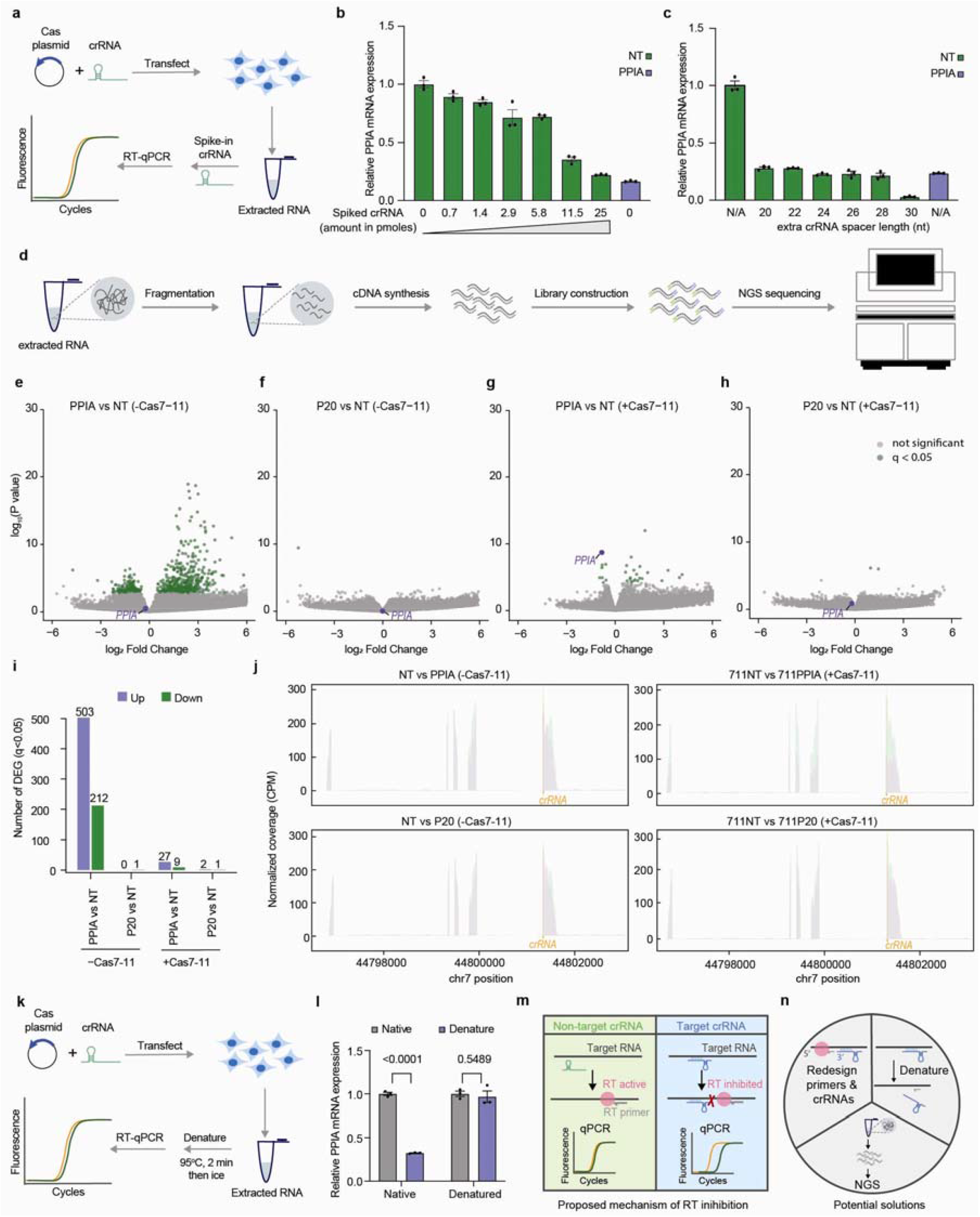
crRNA binding differentially affects RT-qPCR and RNA sequencing. **(a)** Schematic of the post-extraction RNA spike-in assay for RT-qPCR. **(b, c)** Effects of crRNA amount **(b)** and spacer truncation **(c)** on apparent target RNA reduction in the RNA spike-in assay. **(d)** Volcano plots of differential gene expression comparing PPIA crRNA **(e)**, P20 crRNA **(f)**, PPIA crRNA + evoCas7-11 **(g)**, and P20 crRNA + evoCas7-11 **(h)** with the non-targeting control. Differential expression analysis was performed using DESeq2 with a two-sided Wald test and Benjamini–Hochberg correction. Differentially expressed genes were defined as those with an adjusted P value (padj) < 0.05. n = 3 biologically independent samples. **(i)** Numbers of significantly upregulated and downregulated genes. Differentially expressed genes were defined as those with an adjusted P value (padj) < 0.05 and |log⊡ fold change| > 1. **(j)** RNA-seq read coverage surrounding the crRNA target site.**(k)** Schematic of the RT-qPCR workflow incorporating an RNA denaturation step before reverse transcription. **(l)** RNA denaturation before reverse transcription restores accurate RT-qPCR quantification. Statistical significance was determined by ordinary two-way ANOVA with Šídák’s multiple-comparisons test. **(m)** Proposed model illustrating how target-bound crRNAs interfere with reverse transcription, resulting in an apparent reduction in target RNA measured by RT-qPCR. **(n)** Proposed strategies to mitigate crRNA-induced RT-qPCR bias, including primer redesign, RNA denaturation before reverse transcription, and RNA sequencing–based quantification. **(b, c, l)** Data are presented as mean ± s.e.m. from three biologically independent experiments (n = 3).

Although the RNA spike-in assay established that crRNA alone was sufficient to reproduce the apparent reduction in target RNA, even the highest crRNA input tested in vitro failed to match the magnitude of the effect observed following cellular transfection. However, the spike-in assay could not fully explain the stronger apparent reduction in target RNA observed in cells. We therefore sought to determine whether crRNA expression alone genuinely reduced target RNA abundance under cellular conditions by performing RNA sequencing (Fig. 2d). While evoCas7-11 induced robust depletion of the target transcript, cells expressing crRNA alone showed no detectable reduction in target RNA abundance despite exhibiting substantial apparent reduction in target RNA by RT-qPCR in the same samples (Fig. 2e-h). These results demonstrate that crRNA expression alone does not reduce target RNA abundance in cells, indicating that the apparent reduction in target RNA observed by RT-qPCR does not reflect genuine transcript depletion.

Despite having no measurable effect on target RNA abundance, crRNA expression alone induced widespread transcriptomic changes relative to untreated controls (Fig. 2i). Spacer truncation markedly reduced these perturbations, indicating that guide–target complementarity strongly influences off-target transcriptomic responses. This pattern closely paralleled our previous observations with the ΨDNA guide^11^, which likewise produced extensive transcriptomic changes in the absence of its cognate nuclease but substantially fewer upon nuclease co-expression, suggesting that association with the cognate effector limits guide-associated perturbations.

Because RNA-seq failed to detect target depletion despite the apparent RT-qPCR knockdown, we next examined whether crRNA binding biased sequencing coverage across the target transcript. Despite the substantial apparent reduction in target RNA detected by RT-qPCR, RNA-seq read coverage remained uniform across the crRNA-binding site, with no evidence of local depletion or sequencing bias (Fig. 2j). These results demonstrate that crRNA binding does not interfere with RNA sequencing and indicate that the discrepancy between RT-qPCR and RNA-seq arises from the RT-qPCR workflow rather than RNA abundance.

Having established that the apparent reduction in target RNA arose during RNA quantification, we next asked whether it could be eliminated by disrupting crRNA-target interactions before reverse transcription. Heat denaturation of total RNA immediately before reverse transcription completely abolished the apparent reduction in target RNA, restoring RT-qPCR measurements to levels consistent with RNA sequencing (Fig. 2k, l). These results identify reverse transcription as the source of the apparent reduction in target RNA rather than genuine changes in RNA abundance. Together, our results establish RNA denaturation as a simple and readily implementable strategy for accurate RT-qPCR quantification in RNA-targeting CRISPR experiments.

Together, our findings reveal an unrecognized limitation of RT-qPCR-based quantification in RNA-targeting CRISPR studies and have important implications for the interpretation of guide RNA-mediated knockdown experiments. Although the precise molecular mechanism remains to be fully established, our data are consistent with a model in which stable crRNA-target duplexes impede reverse transcription, thereby producing artificially reduced RT-qPCR signals (Fig. 2m). The observation that this artifact is shared across multiple RNA-targeting CRISPR platforms suggests that it represents a general challenge for guide RNA-based RNA targeting technologies rather than a platform-specific phenomenon.

Our findings also have several practical implications (Fig. 2n). Where feasible, RT-qPCR amplicons should preferentially be positioned upstream (5′) of the guide-binding site so that reverse transcription is not required to traverse the crRNA–target duplex. Alternatively, a simple RNA denaturation step before reverse transcription provides a rapid and broadly applicable solution without requiring guide or primer to redesign. For applications requiring transcriptome-wide or orthogonal validation, RNA sequencing offers an unbiased assessment of RNA abundance.

Beyond improving transcript quantification, our transcriptomic analyses also provide guidance for guide RNA design and delivery. Guide RNAs expressed in the absence of their cognate effector induced widespread transcriptomic perturbations, whereas association with the cognate nuclease substantially reduced these effects. These observations suggest that delivery strategies minimizing the accumulation of free guide RNAs, including RNP-based approaches, may improve transcriptomic specificity. Likewise, spacer truncation markedly reduced guide-associated perturbations, suggesting that guide RNAs should be designed to be as short as functionally permissible while maintaining sufficient on-target activity. More broadly, our findings highlight that RNA binding and RNA abundance are not necessarily equivalent in reverse transcription-based measurements, an important consideration for the continued development, evaluation and optimization of RNA-targeting technologies.

## Acknowledgements

We thank Dr. David Corey (University of Texas Southwestern Medical Center) and Dr. Mingyi Xie (University of Florida) for generously providing the AGO knockout cell lines used in this study. We also thank the University of Florida Interdisciplinary Center for Biotechnology Research (UF ICBR) Next-Generation DNA Sequencing Core (RRID: SCR_019152) for sequencing support. This work was supported by the University of Florida, the UF Herbert Wertheim College of Engineering, the Exxon Mobil Gator Alumni Faculty Endowment Funds, NIH-NIAID (R61AI181016), and NIH-NIGMS (R35GM147788). The funding agencies had no role in study design, data collection and analysis, decision to publish, or preparation of the manuscript.

