## Supplementary Information for "Guide RNA binding induces reverse transcription bias during RT-qPCR analysis of RNA-targeting CRISPR/Cas systems"

**The PDF file includes:**

Materials and Methods
Tables S1 to S2
References

**Materials and Methods**

Plasmid Construction

The plasmid encoding evoCas7-11 was generated by introducing the previously reported mutations into the DisCas7-11 plasmid (Addgene #172507) using site-directed mutagenesis. The AsCas12a expression plasmid was described previously[11]. All plasmid constructs were verified by Sanger sequencing.

Oligonucleotide preparation

All single-stranded DNA and RNA oligos including guide RNA, guide DNA, target activators, primers, and fluorescent reporters were obtained from Integrated DNA Technologies (IDT) and diluted in 1x TE Buffer (10 mM Tris, 0.1 mM EDTA, pH 7.5).

Mammalian cell culture

HEK293T (ATCC #CRL-3216) cells were cultured in DMEM high glucose GlutaMAX™ supplement pyruvate (Gibco #10569010), 10% Fetal Bovine Serum (Gibco #A3160902) and 1X Penicillin-Streptomycin. Cells were incubated at 37°C and 5% $CO_2$.

HCT116 wild-type cells and HCT116-derived AGO2-knockout and AGO1/2/3 triple-knockout cell lines were cultured in HyClone™ McCoy's 5A medium (Cytiva, SH30200.01) supplemented with 10% fetal bovine serum (Gibco, A3160902) and 1× penicillin–streptomycin. Cells were maintained at 37 °C in a humidified incubator with 5% $CO_2$.

HCT116-derived AGO2-knockout and AGO1/2/3 triple-knockout cell lines were generously provided by Dr. David Corey (University of Texas Southwestern Medical Center) through Dr. Mingyi Xie (University of Florida).

Mammalian cell transfection

For plasmid and crRNA transfection experiments, HEK293T cells were seeded at 0.2 M cells in a well of 48-well plate 24 h before transfection. Plasmid DNA and synthetic crRNAs were co-transfected using TransIT-X2® transfection reagent (Mirus Bio, MIR6000) according to the manufacturer's instructions. For all transfections, 350 ng of the indicated Cas expression plasmid and the indicated amount of crRNA were mixed with 2 µL of TransIT-X2® in 50 µL of Opti-MEM™ Reduced Serum Medium (Gibco, 31985062). Complexes were incubated for 15 min at room temperature before being added dropwise to pre-seeded 48-well plates.

For the experiments shown in Fig. 1a, 350 ng of evoCas7-11 or AsCas12a expression plasmid was co-transfected with 12.5 nM evoCas7-11 crRNA. For the crRNA titration experiments (Fig. 1c, d), 350 ng of evoCas7-11 expression plasmid was co-transfected with the indicated concentrations of crRNA, whereas crRNA-only groups received the corresponding amount of crRNA without a Cas expression plasmid. For the guide architecture, spacer truncation and Ago-knockout experiments (Fig. 1e-g), 350 ng of the indicated Cas expression plasmid was co-transfected with 12.5 nM guide RNA, whereas guide-only groups received 12.5 nM guide RNA in the absence of a Cas expression plasmid. Unless otherwise indicated, cells were harvested 18 h after transfection for RNA extraction and RT-qPCR analysis.

RT-qPCR for relative quantification of mCherry and endogenous mRNA

Total RNA of samples was extracted using Monarch Total RNA Miniprep Kit (NEB #T2010S) as per manufacturer's instructions. Extracted RNA was later added to the RT-qPCR mix TaqMan™ Fast Virus 1-Step Master Mix (Thermo #4444434). The reaction was performed on an Applied Biosciences QuantStudio™ 5 Real-Time PCR System and multiplexed with FAM probes for target genes, and Cy5 probe for GAPDH as the housekeeping gene. Primers and probes of target (PPIA and RPL4) were ordered from Thermo Fisher. Fold-changes were calculated by ddCt method. One-way or two-way ANOVA with multiple comparison correction was performed to assess the statistical significance of transcript changes, using Prism 10.

Post-extraction RNA spike-in assay

Total RNA was extracted from untreated HEK293T cells using the Monarch Total RNA Miniprep Kit (NEB, T2010S) according to the manufacturer's instructions. For concentration-dependent experiments, 40 ng of purified total RNA was mixed with the indicated amounts of synthetic crRNA and incubated at 37 °C for 30 min before reverse transcription.

For spacer truncation experiments, 40 ng of total RNA was incubated with 25 pmol of synthetic crRNAs containing the indicated spacer lengths under the same conditions. Reverse transcription and RT-qPCR were subsequently performed as described above.

RNA denaturation assay

Following total RNA extraction, RNA samples were heated at 95 °C for 2 min and immediately cooled on ice before reverse transcription. Untreated RNA samples were processed in parallel as native controls. Reverse transcription and RT-qPCR were then performed as described above.

mRNA-Seq

For RNA-seq experiments, HEK293T cells were seeded at $2x10^6$ cells per well in 6-well plates 24 h before transfection. Cells were transfected with either 1 µg of evoCas7-11 expression plasmid together with 50 nM crRNA or 50 nM crRNA alone (in the absence of an evoCas7-11 expression plasmid) using TransIT-X2® transfection reagent (Mirus Bio, MIR6000). Transfection complexes were prepared in 250 µL of Opti-MEM™ Reduced Serum Medium (Gibco, 31985062) according to the manufacturer's instructions and added dropwise to each well. Cells were harvested 18 h after transfection for RNA extraction and RNA sequencing.

After GFP positive cells were sorted, total RNA was extracted from them using Monarch Total RNA Miniprep Kit (NEB #T2010S) as per manufacturer's instructions. RNA integrity (RIN) score of the samples was measured with a QIAxcel Advance with an QIAxcel RNA QC Kit v2.0 (Qiagen #929104) cartridge to proceed with RNA library preparation for Illumina NGS sequencing,

Libraries were prepared with NEBNext® Ultra™ II Directional RNA Library Prep (NEB #E7765S) in conjunction with NEBNext Poly(A) mRNA Magnetic Isolation Module (NEB #E7490) to isolate only mRNA from the samples. Indexes used were NEBNext® Multiplex Oligos for Illumina® Set 1, 2, and 3 (NEB #E7335S, #E7500S, #E7710S). Final libraries were loaded in a QIAxcel DNA Screening Kit (Qiagen #929004) to check for correct size distribution before sequencing and quantifies using Qubit Flex and qPCR.

Total RNA was extracted using the Monarch Total RNA Miniprep Kit (NEB, T2010S) according to the manufacturer's instructions. mRNA was isolated using the Dynabeads™ mRNA Purification Kit (Thermo Fisher Scientific, 61006). Sequencing libraries were prepared using the NEBNext® Ultra™ II Directional RNA Library Prep Kit for Illumina (NEB, E7765S) according to the manufacturer's instructions. Libraries were pooled and sequenced on an Illumina NovaSeq X Plus platform using a NovaSeq X Series 1.5B Reagent Kit (100 Cycle) (Illumina, 20104703) to generate 2 × 150 bp paired-end reads, yielding approximately 30 million reads per sample.

Output pair-end reads from sequencing were aligned with HISAT2 and then converted to sorted bam files with samtools. Then these files were processed using featureCounts to get a count matrix of each gene. Finally, the matrices were used as input for DESeq2 to quantify off-target effects in the transcriptome. Volcano plots were created using R. All computational work was performed on University of Florida's HPC HiperGator.

**Table S1.** Protein amino acid sequences

| Protein | Amino Acid Sequence |
|---|---|
| AsCas12a | MTQFEGFTNLYQVSKTLRFELIPQGKTLKHIQEQGFIEEDKARNDHYKELKPIIDRI<br>YKTYADQCLQLVQLDWENLSAAIDSYRKEKTEETRNALIEEQATYRNAIHDYFIGR<br>TDNLTDAINKRHAEIYKGLFKAELFNGKVLKQLGTVTTTEHENALLRSFDKFTTYF<br>SGFYENRKNVFSAEDISTAIPHRIVQDNFPKFKENCHIFTRLITAVPSLREHFENVK<br>KAIGIFVSTSIEEVFSFPFYNQLLTQTQIDLYNQLLGGISREAGTEKIKGLNEVLNLAI<br>QKNDETAHIIASLPHRFIPLFKQILSDRNTLSFILEEFKSDEEVIQSFCKYKTLLRNE<br>NVLETAEALFNELNSIDLTHIFISHKKLETISSALCDHWDTLRNALYERRISELTGKI<br>TKSAKEKVQRSLKHEDINLQEIISAAGKELSEAFKQKTSEILSHAHAALDQPLPTTL<br>KKQEEKEILKSQLDSLLGLYHLLDWFAVDESNEVDPEFSARLTGIKLEMEPSLSFY<br>NKARNYATKKPYSVEKFKLNFQMPTLASGWDVNKEKNNGAILFVKNGLYYLGIM<br>PKQKGRYKALSFEPTEKTSEGFDKMYYDYFPDAAKMIPKCSTQLKAVTAHFQTH<br>TTPILLSNNFIEPLEITKEIYDLNNPEKEPKKFQTAYAKKTGDQKGYREALCKWIDF<br>TRDFLSKYTKTTSIDLSSLRPSSQYKDLGEYYAELNPLLYHISFQRIAEKEIMDAVE<br>TGKLYLFQIYNKDFAKGHHGKPNLHTLYWTGLFSPENLAKTSIKLNGQAELFYRP<br>KSRMKRMAHRLGEKMLNKKLKDQKTPIPDTLYQELYDYVNHRLSHDLSDEARAL<br>LPNVITKEVSHEIIKDRRFTSDKFFFHVPITLNYQAANSPSKFNQRVNAYLKEHPET<br>PIIGIDRGERNLIYITVIDSTGKILEQRSLNTIQQFDYQKKLDNREKERVAARQAWS<br>VVGTIKDLKQGYLSQVIHEIVDLMIHYQAVVVLENLNFGFKSKRTGIAEKAVYQQF<br>EKMLIDKLNCLVLKDYPAEKVGGVLNPYQLTDQFTSFAKMGTQSGFLFYVPAPYT<br>SKIDPLTGFVDPFVWKTIKNHESRKHFLEGFDFLHYDVKTGDFILHFKMNRNLSF<br>QRGLPGFMPAWDIVFEKNETQFDAKGTPFIAGKRIVPVIENHRFTGRYRDLYPAN<br>ELIALLEEKGIVFRDGSNILPKLLENDDSHAIDTMVALIRSVLQMRNSNAATGEDYI<br>NSPVRDLNGVCFDSRFQNPEWPMDADANGAYHIALKGQLLLNHLKESKDLKLQN<br>GISNQDWLAYIQELRN |
| evoCas7-11[3, 7] | MTTTMKISIEFLEPFRMTKWQESTRRNKNNKEFVRGQAFARWHRNKKDNTKGR<br>PYITGTLLRSAVIRSAENLLTLSDGKISEKTCCPGKFDTEDKDRLLQLRQRSTLRW<br>TDKNPCPDNAETYCPFCELLGRSGNDGKKAEKKDWRFRIHFGNLSLPGKPDFD<br>GPKAIGSQRVLNRVDFKSGKAHDFFKAYEVDHTRFPRFEGEITIDNKVSAEARKL<br>LCDSLKFTDRLCGALCVIRFDEYTPAADSGKQTENVQAEPNANLAEKTAEQIISIL<br>DDNKKTEYTRLLADAIRSLRRSSKLVAGLPKDHDGKDDHKLWDIGKKKKDENSVT<br>IRQILTTSADTKELKNAGKWREFCEKLGEALYLKSKDMSGGLKITRRILGDAEFHG<br>KPDRLEKSRSVSIGSVLKETVVCGELVAKTPFFFGAIDEDAKQTDLQVLLTPDNKY<br>RLPRSAVRGILRRDLQTYFDSPCNAELGGRPCMCKTCRIMRGITVMDARSEYNA<br>PPEIRHRTRINPFTGTVAEGALFNMEVAPEGIVFPFQLRYRGSEDGLPDALKTVLK<br>WWAEGQAFMSGAASTGKGRFRMENAKYETLDLSDENQRNDYLKNWGWRDEK<br>GLEELKKRLNSGLPEPGNYRDPKWHEINVSIEMASPFINGDPIRAAVDKRGTDVV<br>TFVKYKAEGEEAKPVCAYKAESFRGVIRSAVARIHMEDGVPLTELTHSDCECLLC<br>QIFGSEYEAGKIRFEDLVFESDPEPVTFDHVAIDRFTGGAADKKKFDDSPLPGSP<br>ARPLMLKGSFWIRRDVLEDEEYCKALGKALADVNNGLYPLGGKSAIGYGQVKSL |

| | |
|---|---|
| | GIKGDDKRISRLMNPAFDETDVAVPEKPKTDAEVRIEAEKVYYPHYFVEPHKKVE<br>REEKPCGHQKFHEGRLTGKIRCKLITKTPLIVPDTSNDDFFRPADKEARKEKDEY<br>HKSYAFFRLHKQIMIPGSELRGMVSSVYETVTNSCFRIFDETKRLSWRMDAKHQ<br>NVLQKFLPGRVTADGKHIQKFSETARVPFYDKTQKHFDILDEQEIAGEKPVRMWV<br>KRFIKRLSLVDPAKHPQKKQDNKWKRRKEGIATFIEQKNGSYYFNVVTNNGCTSF<br>HLWHKPDNFDQEKLEGIQNGEKLDCWVRDSRYQKAFQEIPENDPDGWECKEG<br>YLHVVGPSKVEFSDKKGDVINNFQGTLPSVPNDWKTIRTNDFKNRKRKNEPVFC<br>CEDDKGNYYTMAKYCETFFFDLKENEEYEIPEKARIKYKELLRVYNNNPQAVPES<br>VFQSRVARENVEKLKSGDLVYFKHNEKYVEDIVPVRISRTVDDRMIGKRMSADLR<br>PCHGDWVEDGDLSALNAYPEKRLLLRHPKGLCPACRLFGTGSYKGRVRFGFAS<br>LENDPEWLIPGKNPGDPFHGGPVMLSLLERPRPTWSIPGSDNKFKVPGRKFYVH<br>HHAWKTIKDGNHPTTGKAIEQSPNNRTVEALAGGNSFSFEIAFENLKEWELGLLI<br>HSLQLEKGLAHKLGMAKSMGFGSVEIDVESVRLRKDWKQWRNGNSEIPNWLGK<br>GFAKLKEWFRDELDFIENLKKLLWFPEGDQAPRVCYPMLRKKDDPNGNSGYEEL<br>KRGEFKKEDRQKKLTTPWTPWA |

**Table S2.** crRNA sequences for endogenous gene silecing and qPCR primers.

| Name | Sequence |
|---|---|
| cr-NT derived from the crRNA in[2] | rG*rU*rU*rGrArUrGrUrCrArCrGrGrArArCrUrCrArCrCrArGrArArGrCrGrUrArCrCrArUrArCrUrCrArCrGrArArC*rA*rG*rC |
| cr-PPIA derived from the crRNA in[15] | rG*rU*rU*rGrArUrGrUrCrArCrGrGrArArCrArArArCrArCrCrArCrArUrGrCrUrUrGrCrCrArUrCrCrArArCrCrA*rC*rU*rC |
| cr-RPL4 derived from the crRNA in[15] | rG*rU*rU*rGrArUrGrUrCrArCrGrGrArArCrArArArCrGrArArGrUrUrCrArGrGrArArCrUrUrCrCrUrCrArArUrArCrGrArU*rG*rA*rC |
| cr-PPIA-28 nt | rG*rU*rU*rGrArUrGrUrCrArCrGrGrArArCrArArArCrArCrCrArCrArUrGrCrUrUrGrCrCrArUrCrCrArArC*rC*rA*rC |
| cr-PPIA-26 nt | rG*rU*rU*rGrArUrGrUrCrArCrGrGrArArCrArArArCrArCrCrArCrArUrGrCrUrUrGrCrCrArUrCrCrA*rA*rC*rC |
| cr-PPIA-24 nt | rG*rU*rU*rGrArUrGrUrCrArCrGrGrArArCrArArArCrArCrCrArCrArUrGrCrUrUrGrCrCrArUrC*rC*rA*rA |
| cr-PPIA-22 nt | rG*rU*rU*rGrArUrGrUrCrArCrGrGrArArCrArArArCrArCrCrArCrArUrGrCrUrUrGrCrC*rA*rU*rC*rC |
| cr-PPIA-20 nt | rG*rU*rU*rGrArUrGrUrCrArCrGrGrArArCrArArArCrArCrCrArCrArUrGrCrUrUrGrC*rC*rA*rU |

| | |
|---|---|
| ΨDNA PPIA | A*A*A*CACCACATGCTTGCCATCCAACCACTCTAGATGTGAATCATCTTT*A*A*T |
| cr-PPIA (13b) | rA*rA*rA*rCrArCrCrArCrArUrGrCrUrUrGrCrCrArUrCrCrArArCrCrArCrUrCrGrUrUrGrUrGrGrArArGrGrUrCrCrArGrUrUrUrGrArGrGrGrGrCrUrArUrUrArC*rA*rA*rC |
| cr-PPIA (13d) | rG*rU*rU*rUrUrArGrUrCrCrCrCrUrUrCrGrUrUrUrUrGrGrGrGrUrArGrUrCrUrArArArUrCrArArArCrArCrCrArCrArUrGrCrUrUrGrCrCrArUrCrCrArArCrCrA*rC*rU*rC |
| qPCR PPIA | Hs99999904_m1 |
| qPCR RPL4 | Hs00973287_g1 |
| qPCR GAPDH For[16] | GCTCCCTCTTTCTTTGCAGCAAT |
| qPCR GAPDH Rev[16] | TACCATGAGTCCTTCCACGATAC |
| qPCR GAPDH Cy5 Probe[16] | Cy5/TCCTGCACC/TAO/ACCAACTGCTTAGCACC/Iowa Black RQ |

*signifies phosphorothioated bonds
r signifies RNA bases
